# Synthetically designed anti-defense proteins overcome barriers to bacterial transformation and phage infection

**DOI:** 10.1101/2025.09.01.673470

**Authors:** Jeremy Garb, David W. Adams, Eliane Hadas Yardeni, Melanie Blokesch, Rotem Sorek

**Affiliations:** Department of Molecular Genetics, Weizmann Institute of Science, Rehovot, Israel; Laboratory of Molecular Microbiology, Global Health Institute, School of Life Sciences, Station 19, EPFL-SV-UPBLO, Ecole Polytechnique Fédérale de Lausanne (EPFL), Lausanne, Switzerland; Department of Life Sciences Core Facilities, Weizmann Institute of Science, Rehovot, Israel

## Abstract

Bacterial defense systems present considerable barriers to both phage infection and plasmid transformation. These systems target mobile genetic elements, limiting the efficacy of bacteriophage-based therapies and restricting genetic engineering applications. Here, we employ a de-novo protein design approach to generate proteins that bind and inhibit bacterial defense systems. We show that our synthetically designed proteins block defense, and that phages engineered to encode the synthetic proteins can replicate in cells that express the respective defense system. We further demonstrate that a single phage could be engineered with multiple anti-defense proteins, yielding improved infectivity in bacterial strains carrying multiple defense systems. Finally, we show that plasmids that express synthetic anti-defense proteins can be introduced into bacteria that naturally restrict plasmid transformation. Our approach can broaden host ranges of therapeutic phages and can improve genetic engineering efficiency in strains that are typically difficult to transform.

## Introduction

Studies from recent years have substantially expanded knowledge on bacterial defense against mobile genetic elements^1^. Whereas restriction enzymes and CRISPR-Cas were traditionally considered the main facets of the bacterial immune system, it is now understood that bacteria encode over 150 different mechanisms to resist invading phages and plasmids^2–6^. An average bacterial genome encodes ∼6 defense systems, with some genomes containing more than 50 systems^7,8^. There is a substantial variation in the sets of defense systems carried by individual bacteria, even among closely related strains of the same species^7–9^.

The newly realized presence of defense systems in bacteria may provide an explanation for the limited success of using phages as therapeutic agents against pathogenic bacteria^10,11^. The process of developing phages as antibacterial compounds usually involves isolation of phages that can infect a set of laboratory-grown strains of a given pathogen^12^. However, as strains of the same species show high variability in the sets of defense systems they encode^7–9^, phages isolated on laboratory strains can be inefficient when infecting in-patient bacteria that carry defense systems different than those present in the laboratory strains. Indeed, phages that efficiently targeted laboratory strains of pathogenic bacteria sometimes failed when deployed in patients infected with the respective pathogen^13^, and it was shown that in some cases the patient-residing strains were resistant to the phage ^14^.

Bacterial defense systems also explain the difficulties in genetic engineering of wild bacteria^15–18^. Genetic engineering of bacteria isolated from the wild is notoriously difficult, and is usually limited to a small set of laboratory-domesticated bacteria defined as “model organisms” that allow DNA transformation^19^. Recent studies reported bacterial defense systems that specifically target plasmids, including Wadjet^20^, prokaryotic Argonautes (pAgo)^17,18^, and Lamassu^21^, and it was shown that these systems can be responsible for the difficulties in maintaining transformed plasmids in bacteria^16,22^. This limitation in the ability to genetically engineer bacteria obstructs the development of bacterial strains for research, industrial, and therapeutic purposes.

With recent progress in the protein design field, it has become possible to design completely synthetic proteins with desired properties^23^. Techniques like RFdiffusion^24^, MaSIF-seed^25^ and BindCraft^26^ now allow de-novo design of proteins that would bind target proteins of choice. In the current study we employ a de-novo protein design approach to engineer synthetic proteins that bind and inhibit bacterial defense systems. We show that such engineered anti-defense proteins can be designed to target functional sites of defense system proteins and inhibit their activities. By engineering phages to express synthetic anti-defense proteins, we show that these proteins allow phages to successfully infect bacteria even if these express multiple defense systems. Additionally, we show that our approach enables the design of plasmids that can be maintained in strains considered hard to transform. This approach holds promise for improving the therapeutic potential of phages, and also for advancing genetic engineering applications by overcoming the intrinsic barriers posed by bacterial defense systems.

## Results

### Synthetically designed proteins that inhibit Thoeris

To examine the feasibility of designing protein inhibitors for bacterial defense systems we selected the Type I Thoeris system from *Bacillus cereus* MSX-D12, a defense system whose mechanism is well characterized^2,27^, as an initial target for inhibition. This system comprises two genes, ThsA and ThsB (Fig. 1a). ThsB senses phage invasion and produces the signaling molecule 3’cADPR (also called 1”-3’ gcADPR), which binds the ThsA protein. Once activated by 3’cADPR, ThsA cleaves the essential molecule NAD^+^ and depletes it from the infected cell^28,29^. To activate ThsA, 3’cADPR binds a specific domain in the ThsA protein called SLOG^27,30,31^. We hypothesized that designing small proteins that bind the molecule-sensing site in the SLOG domain would disrupt its ability to perceive the signaling molecules, thereby impairing the protective function of the system (Fig. 1b).

**Figure 1.**
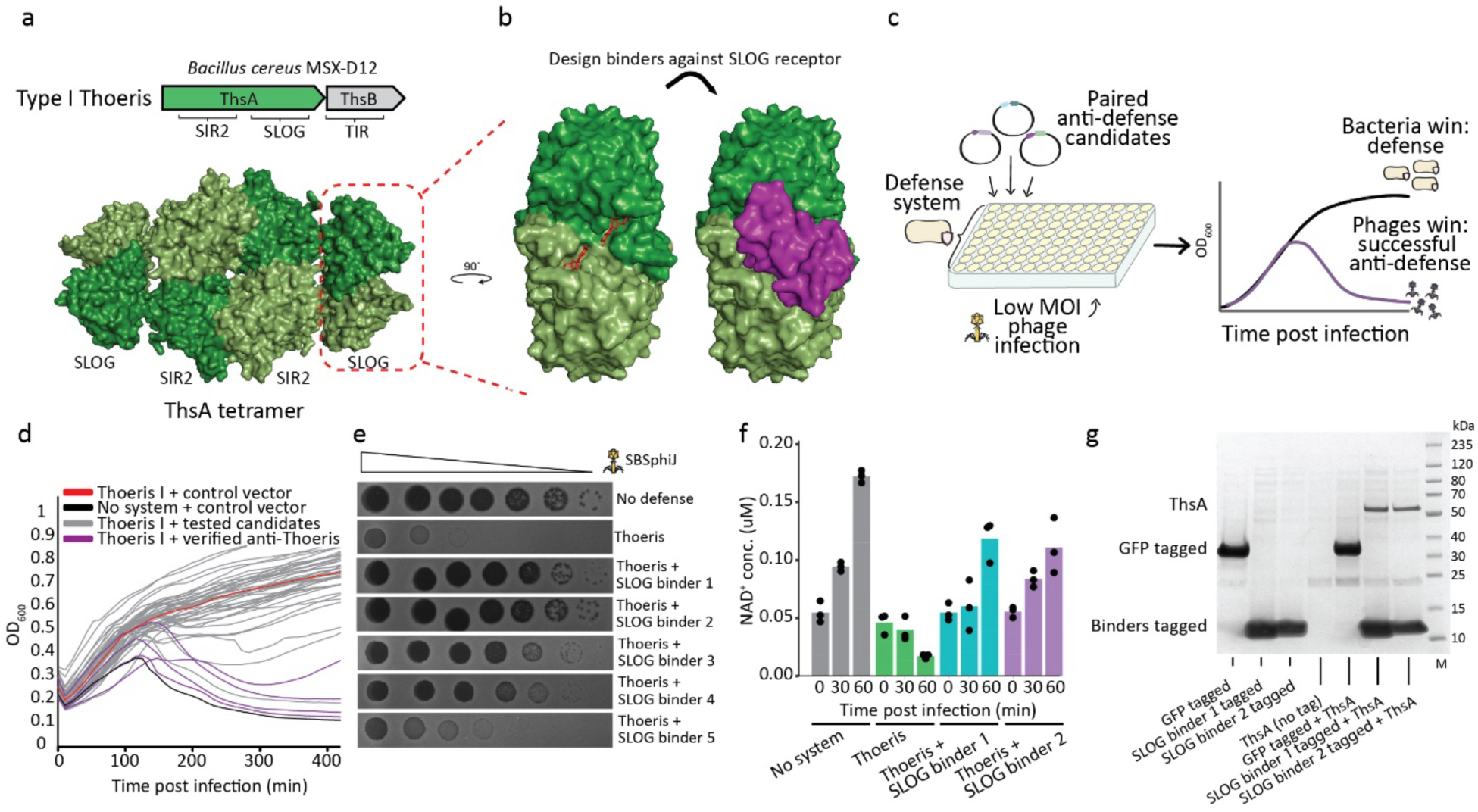
Synthetically designed proteins that inhibit Thoeris. (a) Top, the Thoeris system of *B. cereus* MSX-D12 used in this study. Protein domains are indicated. Bottom, surface representation of the ThsA tetramer (AF3 model). (b) Left, surface representation of the SLOG receptor dimer in complex with 3’cADPR (red molecules)(PDB ID 7UXS)^29^. Right, AF3 model of SLOG binder 2 interacting with the SLOG domain pocket (ipTM=0.84). (c) Schematic representation of the experimental set up. Liquid cultures of bacteria harboring the defense system are grown in a 96 well plate format. Each well is transformed with a different plasmid that expresses a pair of anti-defense candidates, with antibiotics selection added to select for cells that integrated the plasmid. Bacteria are subsequently infected with phages in a low MOI, and OD is measured to identify successful phage infections. (d) Cells expressing the type I Thoeris system from *B. cereus* MSX-D12 were infected with phage SBSphiJ at MOI 0.01. Each curve represents growth of bacteria transformed with a pair of candidate anti-defense genes, with negative control cells transformed with GFP instead (black). Purple curves represent cases where the transformed plasmid reproducibly allowed the phage to propagate and cause eventual culture collapse, suggesting that at least one of the genes in the transformed plasmid inhibited Thoeris defense. (e) Five identified binder proteins capable of overcoming type I Thoeris defense. Shown are tenfold serial dilution plaque assays, comparing the plating efficiency of phage SBSphiJ on bacteria that express the type I Thoeris alone or with a synthetic binder. Images are representative of three replicates. (f) Concentrations of NAD^+^ in cell lysates extracted from SBSphiJ-infected cells as measured by a biochemical NAD^+^ assay. The *x* axis represents minutes post infection, with zero representing non-infected cells. Cells were infected at an MOI of 10 at 30 °C. Bar graphs represent the average of three biological repeats, with individual data points overlaid. (g) Pulldown assays of anti-Thoeris (SLOG targeting) binders and the ThsA complex. Two anti-Thoeris binders and control GFP were C-terminally Twin-Strep tagged, mixed with lysates from ThsA-expressing cells, and pulled down. Shown is an SDS–PAGE gel. Rightmost lane is a size marker.

Using RFdiffusion^24^, a deep learning-based protein design tool, we generated a library of in-silico designed binder backbones aimed at the SLOG domain of the ThsA protein. The SLOG domain in its homodimeric form, and specifically the molecule-binding pocket of the SLOG, was used as a target for binder design (Fig. 1b, Methods). Following backbone design, we used ProteinMPNN^32^ to model sequences for the predicted backbones. Candidate protein designs were co-folded with the SLOG domain using AlphaFold2 (AF2) with target templating and the previously described “AF2 initial guess” feature ^33^, and were scored based on mean predicted aligned error (pAE).

The 96 top scoring candidate protein binders were chosen for experimental validation. We developed a high-throughput screening methodology to assess the efficacy of these candidates in neutralizing Thoeris defense. In this method, pairs of genes encoding candidate proteins are transformed into a *Bacillus subtilis* strain harboring the Thoeris defense system (Fig. 1c). This process takes place in a 96-well plate format where candidate or control plasmids are transformed. The cultures are grown and passaged to fresh media under antibiotics selection, and then the culture is subjected to phage infection in liquid media at low MOI. Under normal conditions, an active Thoeris system protects the host cells, allowing the culture to continue growing in the presence of phage infection. Conversely, interference with Thoeris function by a protein binder results in phage-induced lysis of the culture (Fig. 1c).

Through this screening approach we identified four constructs which, when transformed to Thoeris-expressing cells, demonstrated a capacity to inhibit Thoeris defense (Fig. 1 d). As each of these constructs expressed a pair of candidate binder proteins, we then separated each pair into individual genes and used plaque assays to test the activity of each candidate on its own (Fig. S1). This process identified five genes that inhibited Thoeris when co-expressed with it (Fig. S1, Table S1). In *B*. *subtilis,* Thoeris reduces the plating efficiency of phage SBSphiJ by five orders of magnitude (Fig. 1e). Two of the five inhibitors were able to completely abolish Thoeris defense, generating plating efficiency comparable to the control strain in which Thoeris is absent (Fig. 1e). The three other inhibitors had a partial counter-Thoeris effect (Fig. 1e, S1).

As the designed inhibitory proteins presumably block ThsA from sensing the signaling molecule, it is anticipated that ThsA would not be activated to deplete NAD^+^ during phage infection in the presence of these proteins. We used a biochemical assay to monitor NAD^+^ levels in infected cells at various time points following initial infection. As previously demonstrated^27^, when Thoeris-containing cells are infected by phage, cellular NAD^+^ decreases sharply between 30 and 60 minutes from the onset of infection (Fig. 1f). NAD^+^ levels did not decrease in cells in which our designed binders were co-expressed with Thoeris, confirming that the system is not functional in the presence of the designed proteins (Fig. 1f).

To verify that the synthetic proteins inhibit Thoeris by direct binding to ThsA, we tagged two of the synthetic proteins with a C-terminal Twin-Strep tag^34^, and mixed lysates from cells expressing the tagged proteins with lysates from ThsA-expressing cells. Pulldown assays from the mixed cell lysates showed direct interaction between ThsA and the synthetic proteins (Fig. 1g). We then purified one of these synthetic binders and measured its affinity for binding purified ThsA, yielding a dissociation constant (*K*_d_) of ∼60nM as determined by surface plasmon resonance (SPR) (Fig. S2).

As we capped the size of the designed proteins at 100 amino acids, the five synthetic proteins that inhibit ThsA are all short (49-81 amino acids) and show no sequence similarity to sequences in public databases (Table S1). Examining the AlphaFold-predicted structures of these proteins shows that these are mostly alpha-helical proteins (Figure S3). AlphaFold predictions suggest that each of these proteins binds the SLOG domain in a way that would sterically interfere with the SLOG ability to bind the 3’cADPR signaling molecule (Figure S4).

We next used the same design strategy to engineer candidate proteins that would bind and inhibit the ThsB protein of Thoeris (Fig. 2a). The computational process was targeted to seek proteins that bind either the ThsB active site or regions in predicted protein-protein interaction interfaces. Following experimental testing of candidate designed proteins, this process yielded one protein that was able to disrupt Thoeris defense when co expressed with the system (Fig. 2b, S5). The successful binder targeted a region of ThsB predicted to participate in protein-protein interactions (Fig. 2a, S6).

**Figure 2.**
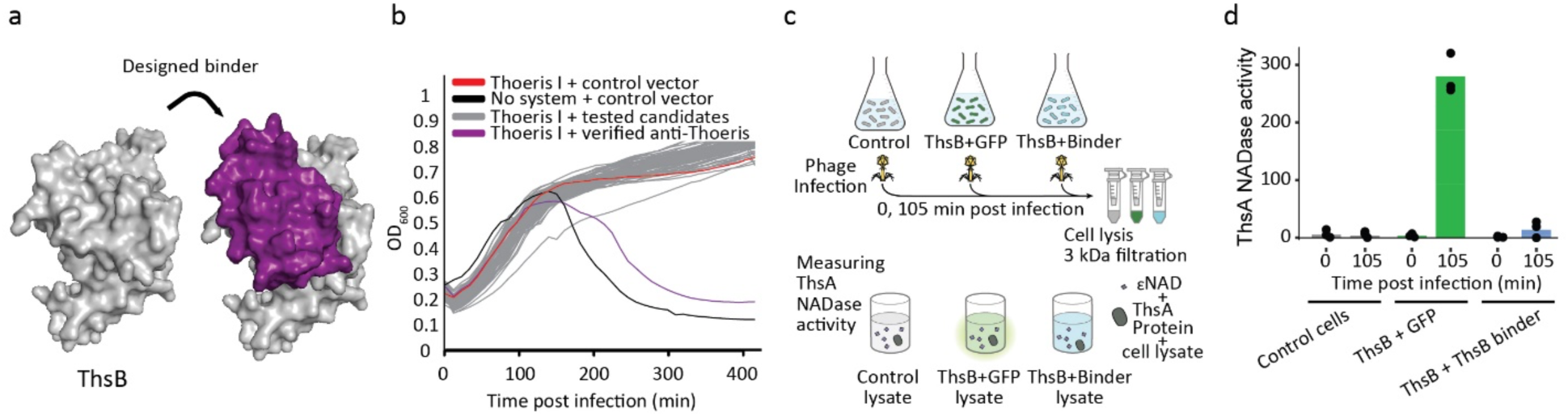
A synthetically designed protein that targets ThsB. (a) Surface representation of the ThsB protein of Thoeris with the ThsB binder, modeled by AF3 (ipTM=0.92). (b) Same experiment as in Fig. 1d but with candidate binders targeting the ThsB protein. (c and d) Binder protein prevents accumulation of 3’cADPR in Thoeris-expressing infected cells. Cells expressing ThsB from *B. cereus* MSX-D12, or co-expressing both ThsB and the anti-ThsB binder, were infected with phage SBSphiJ at a MOI of 10. Cell lysates were extracted before infection (t=0) and 105 mins after infection and were filtered to retain small molecules. Filtered lysates were then incubated with purified ThsA, and the NADase activity of ThsA was measured using a nicotinamide 1,N6-ethenoadenine dinucleotide (εNAD) cleavage fluorescence assay. Y-axis represents Δ εNAD fluorescence per minute. Bars represent the mean of three experiments, with individual data points overlaid. Control cells represent experiments with lysates from cells with no defense system and cells expressing GFP instead of the binder protein.

We expected that binding of the synthetic protein to ThsB would inhibit its ability to produce the 3’cADPR signaling molecule in response to phage infection. To test this hypothesis, we extracted lysates from phage-infected cells that express ThsB and filtered them to retain small molecules (Fig. 2c). As was previously shown^27,28^, lysates from infected cells expressing ThsB triggered the NADase activity of purified ThsA in vitro, confirming that the TIR protein ThsB produces 3’cADPR during infection by phage SBSphiJ (Fig. 2d). In contrast, lysates from infected cells co-expressing ThsB together with the synthetic protein failed to activate ThsA, showing that the synthetic protein inhibited 3’cADPR production in these cells (Fig. 2d).

### Synthetic proteins that inhibit Avs defense

To test if synthetic inhibitors can be designed for defense systems other than Thoeris, we directed our focus toward the Antiviral STAND (Avs) family of bacterial anti-phage proteins^4,35^. These proteins are considered the bacterial ancestors of the eukaryotic NLR immune protein family^36,37^. Similar to eukaryotic NLRs, Avs proteins exhibit a tripartite domain architecture with a central NTPase, an extended C-terminal domain comprising tetratricopeptide (TPR) repeats serving to bind and sense phage proteins, and an N-terminal effector domain that causes cell death or growth arrest once activated^38^. Recognition of a phage protein by the C-terminal domain causes Avs tetramerization and effector activation^35,39^.

We selected Avs1 from *Bacillus cereus* VD102 (Fig. 3a), which, when expressed in *B. subtilis,* confers protection against phage SBSphiJ infection (Fig. S7). The operon architecture of this Avs1 system involves two more genes in addition to the main NLR-like protein, one of them annotated as a hydrolase from the metallo-β-lactamase family, which is hypothesized to participate in the effector activity of the system (Fig. 3a). We selected four possible target sites within the Avs system for binder design: (i) the TPR region of the NLR-like protein, (ii) the ATPase domain of the NLR-like protein, (iii) the protease domain of the NLR-like protein, and (iv) the active site of the Avs-associated hydrolase protein (Fig. 3a).

**Figure 3.**
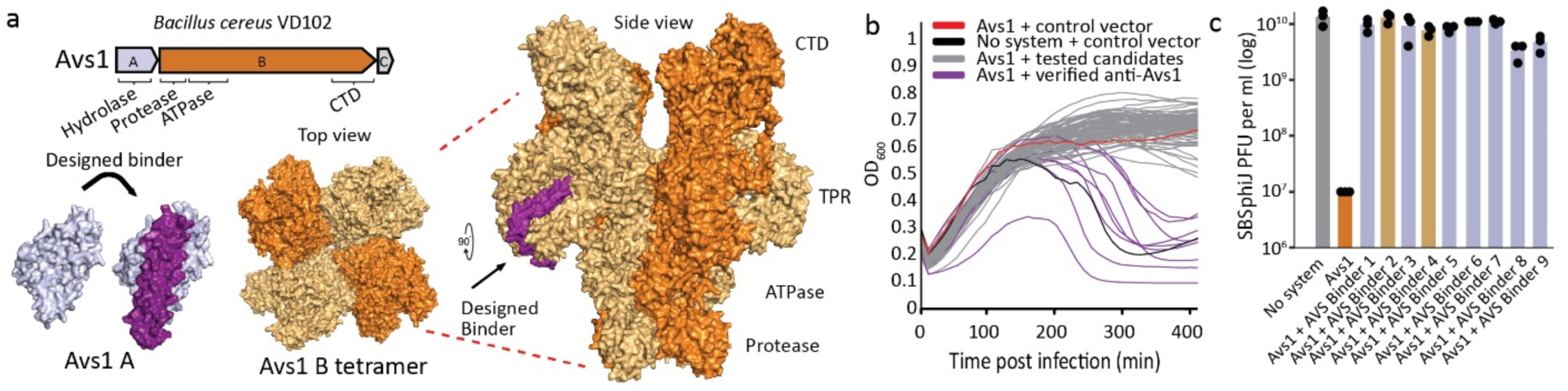
Synthetically designed proteins that inhibit Avs1. (a) Top, the Avs1 system of *B. cereus* VD102 used in this study. Protein domains are indicated. Bottom, surface representation of an AF3 model of Avs1A and Avs1B with example binders (ipTM=0.86 and 0.9, respectively). Due to its size, the Avs1B tetramer was modeled by joining and aligning two AF3 models. (b) Cells expressing the Avs1 system from *B. cereus* VD102 were infected with phage SBSphiJ at MOI 0.01. Each curve represents growth of bacteria transformed with a pair of candidate anti-defense genes, with negative control cells transformed with a plasmid expressing GFP (black). Purple curves represent cases where the transformed plasmid allowed the phage to propagate and cause eventual culture collapse, suggesting that one of the genes in the transformed plasmid inhibited Avs1 defense. (c) Efficiency of plating (EOP) for phage SBSphiJ infecting the control *B. subtilis* BEST7003 strain (no system), *B. subtilis* BEST7003 with Avs1 cloned from *B. cereus* VD102, or cells co expressing Avs1 and single binders. Wheat colored bars represent binders targeting the Avs1B repeat region; purple bars represent Avs1A binders. Data represent plaque-forming units (PFU) per milliliter. Bar graphs depict average of three independent replicates, with individual data points overlaid. Data presented in this panel also appear in Figure S7.

Employing the above-described screening methodology, we identified nine synthetic proteins that facilitated phage infection by overcoming Avs1-mediated defense (Fig. 3b, S7). Two of the synthetic proteins targeted the repeat region of the NLR-like Avs protein and the remaining seven targeted the associated hydrolase (Fig. 3a). When co-expressed with Avs1, these binders completely abolished defense, restoring phage plating efficiency to levels comparable to those observed in cells lacking the Avs1 system (Fig. 3c, S7).

### Engineered phages can overcome multiple defenses

To examine the ability of the designed synthetic proteins to inhibit bacterial defenses also when expressed from phage genomes, we chose the *Bacillus* phage SBSphiJ which is naturally blocked by both Thoeris and Avs1 (Figures 1e, 3c). We engineered individual anti-Thoeris or anti-Avs proteins into the phage genome, at a locus previously validated for having no adverse effect on phage viability ^28^. The synthetic inhibitors were designed under the control of either a natural phage promoter^28^ or a constitutively active bacterial promoter (Fig. 4a, b, Methods). Our data show that a phage engineered to express the anti-Thoeris protein successfully infected Thoeris-expressing bacteria, and similarly, the phage with the synthetic Avs1 inhibitor overcame Avs1 defense (Fig. 4b).

**Figure 4.**
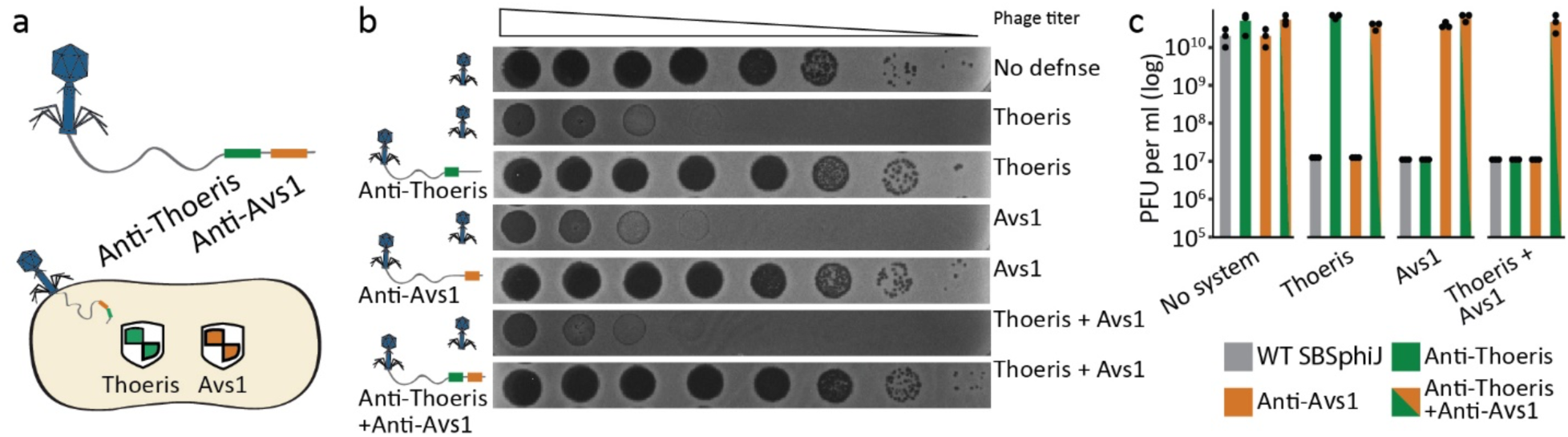
Engineering phages that can overcome multiple defenses. (a) Schematic representation of an engineered phage that carries synthetic proteins designed to overcome multiple defense systems. (b) A SBSphiJ phage carrying binder proteins capable of overcoming Thoeris, Avs1, or both Thoeris and Avs1 defense. Shown are tenfold serial dilution plaque assays, comparing the plating efficiency of phage SBSphiJ on bacteria that express Thoeris alone, Avs1 alone, or both systems. Engineered SBSphiJ phages with individual binders or both binders infect relevant strains. Images are representative of three replicates. (c) Efficiency of plating (EOP) for SBSphiJ phage carrying binder proteins against Thoeris, Avs1, or both, infecting bacteria that express Thoeris alone, Avs1 alone, or both systems. Data represent plaque-forming units (PFU) per milliliter. Bar graphs depict average of three independent replicates, with individual data points overlaid.

Since naturally occurring bacteria often contain more than one defense system, we sought to test whether phages could be engineered with an array of proteins designed to overcome multiple defenses. To this end, we engineered SBSphiJ to express an operon encoding both a synthetic anti-Thoeris protein and an anti-Avs1 protein (Table S2). This phage was able to replicate on *B. subtilis* cells expressing both Thoeris and Avs1 (Fig. 4b, c). As expected, phages engineered with only one of the inhibitor proteins were not able to successfully infect a bacterium in which both systems were co-expressed (Fig. 4c).

To test whether the synthetic proteins inflict burden on phage infectivity, we next performed competition assays between wild type SBSphiJ and the phage strains engineered with the synthetic anti-defense proteins. The synthetically modified phages were not outcompeted by the wild type phage strain when replicating on *B. subtilis* cells expressing no defense system, suggesting that under the experiment conditions, the synthetic proteins do not incur fitness cost on the phage (Fig. S8). The synthetic phages were able to outcompete wild type SBSphiJ when grown on strains expressing either Thoeris, or Avs1, or both (Fig. S8). Together, our data demonstrate that phages can be engineered with synthetically designed defense inhibitors to overcome bacterial defense systems.

### Designed inhibitors against DdmDE rescue plasmids from clearance

Next, we sought to test whether de novo designed anti-defense proteins can allow plasmid transformation into bacterial hosts that are naturally recalcitrant to plasmid maintenance. *Vibrio cholerae* El Tor strains, which are responsible for the ongoing seventh cholera pandemic^40^, are notoriously difficult to stably transform, as incoming plasmids are typically lost after several generations unless strong selective pressure is applied to maintain them^16^. In agreement with this observation, plasmids are naturally absent from the majority of patient-derived isolates^41–43^. It was recently shown that two defense systems actively eliminate plasmids from seventh pandemic *V. cholerae*^16^. These systems comprise the Lamassu-like system DdmABC and the pAgo-like system DdmDE, both encoded on pathogenicity islands specific to pandemic *V. cholerae* strains^16^. The majority of the plasmid clearance phenotype, particularly for small non-conjugative plasmids, is due to the activity of the DdmDE system^16^, and we hence focused on this system for the design of synthetic inhibitors.

The DdmDE system comprises two proteins (Fig. 5a). DdmE is a catalytically inactive pAgo protein that utilizes 5′-phosphorylated DNA guides to recognize plasmid DNA, while DdmD is a helicase-nuclease that naturally exists in the cell in an autoinhibited dimeric state. Upon target recognition by DdmE, DdmD is recruited to the non-target strand, undergoes dimer disassembly, and degrades the plasmid DNA leading to plasmid elimination^16,22,44,45^. We targeted four different sites in the DdmDE system for the design of synthetic binding proteins: (i) the DNA-binding groove of the DdmE PIWI domain, (ii) the DdmE-DdmD interaction interface, (iii) the DNA-binding groove of the DdmD helicase domain, and (iv) the nuclease pocket of DdmD (Fig. 5a). We allowed the computational pipeline to suggest an overall of ∼80,000 candidate binders targeting these sites, and retained 993 top-scoring candidates based on PAE scores from AF2 co-folding predictions.

**Figure 5.**
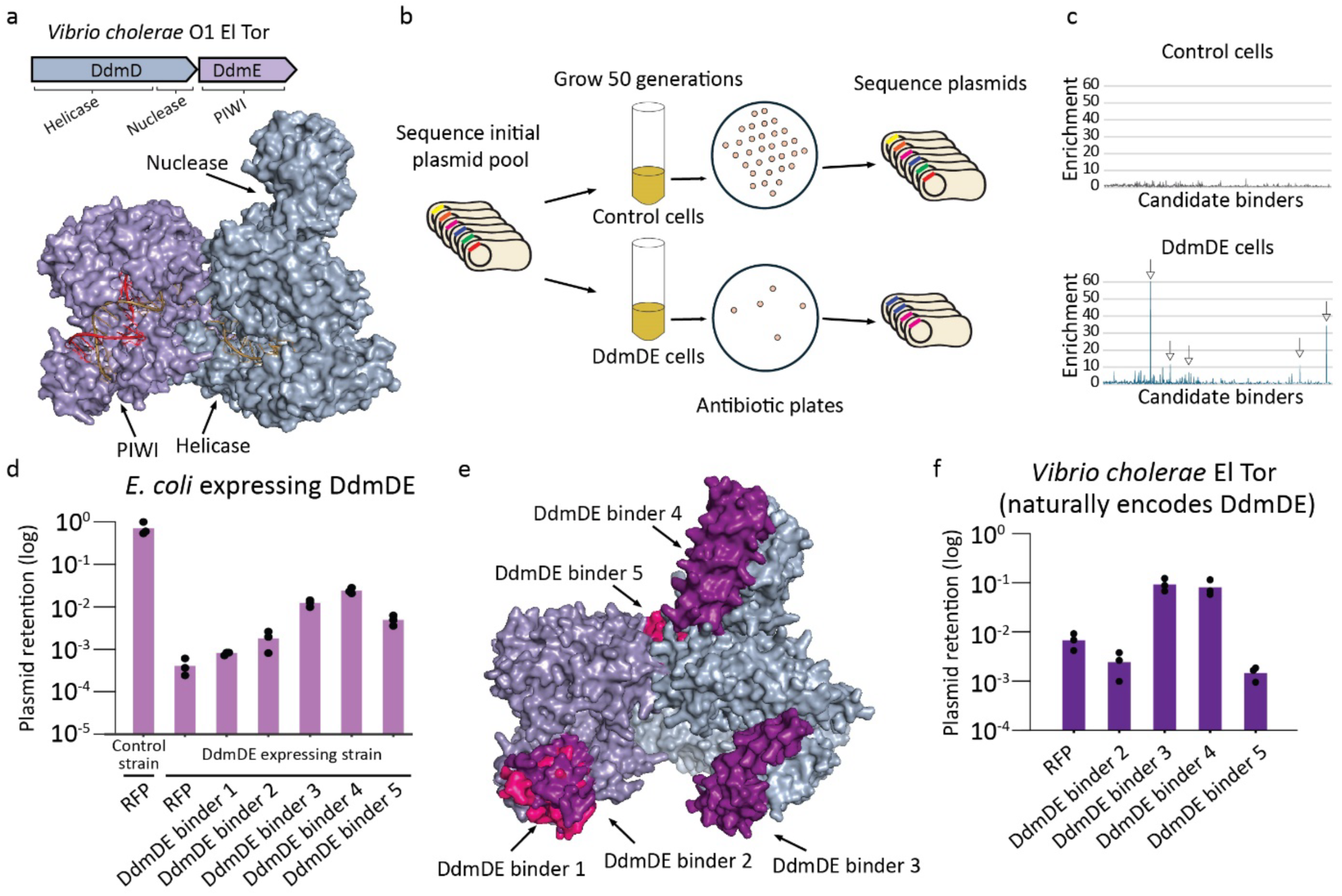
Synthetically designed inhibitors targeting DdmDE rescue plasmids from clearance. (a) Top, the DdmDE system of *V. cholerae* O1 El Tor used in this study. Bottom, surface representation of the DdmDE heterodimer (PDB ID 9EZY)^22^. Domain locations are indicated. Guide DNA (red) and plasmid DNA (yellow) are shown. (b) Schematic representation of the experimental set up. The starting point is a library of *E. coli* bacteria with or without DdmDE, carrying the pool of plasmids, each expressing a single candidate inhibitor. Cultures of these bacteria were grown for 50 generations and plated on antibiotic-containing media. Colonies that grew were collected, and their plasmids were sequenced. (c) Enrichment was calculated by dividing reads per candidate gene in the initial library by the reads per candidate gene at the end point of the experiment. Read counts were normalized to total reads per sample. Top, no enrichment of candidate plasmids in control cells. Bottom, enrichment of candidate plasmids in DdmDE-expressing cells. Shown is a representative of three replicates. All replicates are shown in Fig. S9. (d) Plasmid retention in *E. coli* cells heterologously expressing the *V. cholerae* DdmDE system. Plasmid retention is calculated by dividing CFUs (colony forming units) obtained on antibiotic-containing media by CFUs on non-selective media after 50 generations of growth. Control is an RFP-expressing plasmid. (e) Surface representation of the DdmDE complex modeled with the verified inhibitors. Structures for DdmDE binders 1-4 were modelled via AF3. DdmDE binder 5 was modeled with a cropped section of DdmE and then aligned to the complex structure (Methods) (f) As in panel d, plasmid retention assays were performed in *V. cholerae* O1 El Tor, which naturally encodes the DdmDE system, using either an RFP-expressing control plasmid or plasmids expressing each of the top four binder candidates.

To test whether any of the de novo designed candidates can inhibit DdmDE in vivo, we took advantage of the fact that DdmDE efficiently clears plasmids from transformed cells. We reasoned that plasmids carrying a DdmDE-inhibiting synthetic gene will be retained in DdmDE-expressing cells, while plasmids with non-functional candidates will be lost if transformed to these cells due to DdmDE activity. We therefore adopted a pooled screening approach. The 993 candidate binders were cloned on a plasmid backbone that also encodes an antibiotic selectable marker, and these plasmids were transformed as a pooled library into *E. coli* cells expressing a genome-integrated copy of the DdmDE system from *V. cholerae*. As a control, the same pool was transformed into cells harboring an empty cassette at the genomic integration site (Fig. 5b). Following 50 generations without antibiotics selection, cells from the pooled library were plated on antibiotic-containing LB-agar plates (Fig. 5b). As expected, ∼10,000-fold fewer colonies were observed in the library transformed to DdmDE cells as compared to control cells, due to plasmid elimination by DdmDE. We then sequenced the candidate genes in the plasmids pool extracted from surviving colonies, and sought candidate genes who were significantly enriched in the final pool as compared to their abundance in the initial pool (Fig. 5b). Five genes were reproducibly enriched in final plasmids pools from DdmDE cells, suggesting that these might inhibit DdmDE (Fig 5c, S9). No significant enrichment for any candidate gene was found in cells lacking DdmDE (Fig. 5c).

Each of the five candidate binders was cloned on a plasmid and tested individually for its ability to promote plasmid retention in *E. coli* cells expressing DdmDE. All five candidates showed activity when tested individually, and plasmids carrying candidate binders showed up to a 100-fold elevation in plasmid retention rates in comparison to a control plasmid expressing RFP (Fig. 5d). Incidentally, these binders spanned the four functional sites of DdmDE (Fig. 5e). The synthetic proteins designed to bind the nuclease and helicase domains in DdmD showed the highest activity, validating these sites as targets for DdmDE inhibition (Fig 5d, 5e).

While the above experiments were performed in a heterologous system in which *E. coli* expressed the *V. cholerae* DdmDE, we wished to test whether the de novo designed proteins can promote plasmid retention in native settings. For this, we chose the four binders that showed the most notable anti-DdmDE activity when tested in *E. coli*, and transformed plasmids with each of these binders into a wild type *V. cholerae* O1 El Tor strain. Remarkably, two of these proteins conferred the plasmids with significant resistance against DdmDE-mediated plasmid clearance (Fig 5f). These results demonstrate that our approach is not only effective in a laboratory model but also holds promise for facilitating plasmid retention in a native bacterial strain, highlighting the potential of our approach for domesticating transformation-recalcitrant bacteria.

## Discussion

In this study, we demonstrate that synthetically designed anti-defense proteins can effectively inhibit bacterial defense systems. Engineering these synthetic inhibitors into phage genomes or into plasmid sequences allowed the respective mobile element to overcome defenses by three independent defense systems (Thoeris, Avast, DdmDE) in three different bacteria (*B. subtilis*, *E. coli* and *V. cholerae*). These results suggest a broad applicability for our approach.

In our proof-of-concept study we show that two inhibitors engineered in a single phage can render the phage resistant to two independent defense systems. As the inhibitors we design are short (49-98 amino acids), one can potentially engineer many more such inhibitors in the same phage. Assuming an average size of 200 base pairs per inhibitor, we envision that cassettes of 10 or more inhibitors could be engineered as an operon on a single phage, potentially resulting in phages that resist 10 or more defense systems. Indeed, it was previously shown that phages T7 and Lambda can be engineered with genomic cassettes sized up to ∼2,000 and ∼4,500 bases, respectively^46,47^.

One of the limitations of phage therapy applications is the resistance of patient-resident pathogens to therapeutic phages propagated on laboratory strains of the pathogen, potentially due to different sets of defense systems present in the patient-residing bacterium^13,48^. In a possible future application of our approach, the spectrum of defense systems encoded by the pan-genome of the pathogen can be mapped, and a synthetic inhibitor can be designed for each of these systems. Then, for individual patients, an array of defense inhibitors could be selected based on the presence of the respective defense systems in the patient-resident pathogen. These inhibitors could be engineered into a phage template genome, generating a therapeutic phage that can overcome all defense systems in the target pathogen.

Beyond phage therapy, our findings provide a path for domesticating wild bacteria into model species by allowing genetic manipulation. Many industrially and medically relevant bacterial strains are recalcitrant to transformation due to their defense mechanisms against foreign DNA^49^. Overcoming these transformation barriers has proven highly beneficial in numerous contexts - for instance, enabling the use of non-model organisms in industrial bioconversion processes^50^, harnessing *Streptomyces* species for the production of valuable biological compounds^51^, optimizing *Bacillus* strains for food industry applications^52^, and advancing our understanding of human pathogens such as *Staphylococci* species^53,54^. These examples highlight the broad potential of improved genetic accessibility across diverse bacterial taxa. Notably, species belonging to phyla such as *Aquificae* and *Chloroflexi* do not seem to have any reported cases of transformation in the literature^55,56^. We envision that future use of our approach will facilitate the genetic manipulation of previously intractable bacterial species, broadening the scope of molecular biology applications.

One limitation of the inhibitor design methodology is the relatively low rate of binders that were verified experimentally out of the computationally designed ones. In the current study, we addressed this challenge by testing binders in a semi-high throughput manner, either in a 96-well plate format or in a pooled plasmid approach. Nevertheless, given the low success rates, designing synthetic inhibitors for hundreds of different defense systems would be labor-intensive and cost-ineffective. With the rapid pace of innovations in the field of computational protein design^26^, we envision that future tools would generate synthetic proteins with higher rates of success.

Another limitation of the inhibitors we design is that they likely target the specific proteins against which they were aimed, but not necessarily homologs of these proteins. Naturally occurring viral proteins that inhibit host immune proteins can evolve to bind conserved features in the proteins they target, rendering them active against a broad range of defense system homologs^57^. Future studies could further expand our approach to design synthetic binders that deliberately bind conserved structural features in the target protein. Targeting highly conserved elements may also reduce the likelihood of bacterial escape mutations, as altering these features could compromise the integrity of the defense system itself.

The convergence of synthetic biology, machine learning, and deep mechanistic understanding of bacterial defense systems, now lays the foundation for tailored counter-defense solutions. While our current study focuses on enhancing phage host range and improving plasmid transformation via synthetic protein binders, other applications of synthetic biology in the field of bacterial immunity are imaginable. For example, future studies can tailor protein binders to alter defense system specificities, or to design proteins that sequester immune signaling molecules. The vast design space of synthetic proteins offers a near-limitless reservoir of possibilities.

## Methods

### Bacterial strains

*B. subtilis* strain BEST7003 (obtained from I. Mitsuhiro at Keio University, Japan) was grown in MMB (lysogeny broth (LB) + 0.1 mM MnCl_2_ + 5 mM MgCl_2_, with or without 0.5% agar) at 30 °C. Whenever applicable, media were supplemented with spectinomycin (100 μg mL^−1^) and chloramphenicol (5 μg mL^−1^), to ensure selection of transformed and integrated cells.

*E. coli* strain MG1655 (ATCC 47076) was grown in MMB at 37 °C. Whenever applicable, media were supplemented with ampicillin (100 μg mL^−1^) or chloramphenicol (30 μg mL^−1^) or kanamycin (50 μg mL^−1^), to ensure the maintenance of plasmids.

Type I Thoeris from *Bacillus cereus* MSX-D12 (producing 3’cADPR) was cloned previously under the control of its native promoter to the *amyE* locus of the *B. subtilis* BEST7003 genome^2^. Avs1 from *Bacillus cereus VD102* was cloned similarly under the control of its native promoter to the *amyE* locus.

A1552, the *V. cholerae* O1 El Tor strain used in this work^58^, is a fully sequenced toxigenic O1 El Tor (Inaba) representative of the ongoing seventh cholera pandemic^59^.

For the genomic integration of DdmDE into *E. coli*, as previously described^16^, a mini-Tn*7* transposon carrying *araC* and *DdmDE,* or an empty cassette under the control of the *P*_BAD_ promoter was integrated into a neutral chromosomal locus downstream of *glmS* by triparental mating.

### Phage strains

The *B. subtilis* phage SBSphiJ (Genbank: LT960608.1) isolated by us in a previous study^2^ was used to perform all infection experiments (MOI 0.01, unless stated otherwise).

Engineered SBSphiJ phages used in this paper are listed in Table S2.

Phages were propagated on *B. subtilis* BEST7003 by picking a single phage plaque into a liquid culture grown at 37 °C to an optical density at 600 nm (OD_600_) of 0.3 in MMB broth until culture collapse. The culture was then centrifuged for 10 min at 3,200 *g* and the supernatant was filtered through a 0.2-µm filter to get rid of remaining bacteria and bacterial debris.

### Plasmid construction for use in *V. cholerae*

Fragments encoding candidate genes or RFP under the control of LacI, the LlacO-1 promoter and the *rrnB* T1 terminator, were amplified from the relevant pBbA6c template plasmid and cloned into the *Dra*I site of the *V. cholerae* model plasmid pSa5Y-Kan ^16^. All constructions were verified by PCR and Oxford Nanopore Technologies (ONT)-based full plasmid sequencing (Microsynth AG, Switzerland). Plasmids were propagated in *E. coli* strain TOP10 and introduced into *V. cholerae* by electroporation, as previously described^16^.

### Cloning and transformation of candidate anti-defense genes

Candidate anti-defense genes were codon optimized for *B. subtilis* and synthesized by Twist Bioscience as operonic pairs under the control of a single IPTG promoter. The sequence ‘taataaggaggacaaac’ was added between the two open reading frames to generate a ribosome binding site. These fragments were subsequently added to a Gibson reaction (NEBuilder^®^ HiFi DNA Assembly) with the backbone of the thrC-Phspank vector^28^, which carries a low-copy p15a origin of replication and a chloramphenicol resistance marker. The resulting reaction was then introduced into *Bacillus subtilis* BEST7003 cells, where the respective defense system was integrated into the *amyE* locus.

Transformations were performed using a 96-well format. Prior to transformation, *B. subtilis* BEST7003 cells with or without a defense system were cultured in MC medium at 37 °C for 3 hours, as previously described ^2^. MC medium was composed of 80 mM K2HPO4, 30 mM KH2PO4, 2% glucose, 30 mM trisodium citrate, 22 μg mL^−1^ ferric ammonium citrate, 0.1% casein hydrolysate (CAA) and 0.2% potassium glutamate. Aliquots of 200 µL were transferred to a deep 96-well plate, and 10 µL of the Gibson reaction (NEBuilder^®^ HiFi DNA Assembly) was added to each well. After 3 hour incubation at 37 °C, 1 mL of MMB medium supplemented with spectinomycin (100 μg/mL) and chloramphenicol (5 μg/mL) was added to each well to dilute the MC medium, and the plate was incubated overnight at 30°C. On the following day, a 10 µL aliquot from each well was transferred into 990 µL of fresh MMB containing spectinomycin (100 μg/mL) and chloramphenicol (5 μg/mL) for a single passage. After overnight incubation at 37 °C, the cultures were used as starters for infection experiments in liquid cultures.

### Cloning of candidate anti-defense genes for phage engineering

Candidate anti-defense genes were codon-optimized for *Bacillus subtilis* and synthesized by Twist Bioscience. For constructs containing two synthetic anti-defense genes, the intergenic region included the sequence gggtaaatgtgagcactcacaattcattttgcaaaagttgttgactttatctacaaggtgtggcataatgtgtgtaattgtgagcggataacaattaagcttagtcgacagctagctgattaactaataaggaggacaaac, which introduces a *Phspank* promoter between the two open reading frames. These constructs were cloned into a plasmid backbone previously used for *tad1* knock-in^28^, flanked by ∼1.2 kb upstream and downstream homologous arms corresponding to the SBSphiJ phage genome (primers: F – gaacaactggagggaaaacacttg; R – agctacctcctataggtataaaatttg). This construct drives expression from the native promoter of the anti Thoeris gene *tad1*.

Cloning was performed using the NEBuilder HiFi DNA Assembly kit (NEB, cat # E5520S), and assembled plasmids were transformed into *E. coli* NEB 5-alpha competent cells. Verified plasmids were then transformed into the *thrC* locus of *B. subtilis* BEST7003.

The resulting *B. subtilis* strain, carrying the anti-defense gene(s) and promoter flanked by phage-homologous regions, was infected with SBSphiJ phage at a multiplicity of infection (MOI) of 0.01. Lysates were collected post-infection, and recombinant phages were selected using a Cas13a-gRNA-SBSphiJ targeting strain^60^, plated on agar supplemented with 0.2% xylose to induce Cas13a expression. Resistant phages were plaque-purified three times on *B. subtilis* BEST7003 and verified by PCR and Sanger sequencing for the presence of the candidate anti-defense gene(s).

### Plaque assays

Phage titer was determined using the small drop plaque assay method^61^. A 300 μL of overnight culture of *B. subtilis* was mixed with 30 mL MMB 0.5% agar supplemented with 1mM IPTG and poured into a 10 cm square plate followed by incubation for 1 h at room temperature. Tenfold serial dilutions in MMB were carried out for each of the tested phages and 10 µL drops were put on the bacterial layer. After the drops had dried up, the plates were inverted and incubated overnight at 30 °C. PFUs were determined by counting the formed plaques after overnight incubation and lysate titer was determined by calculating PFUs per milliliter. When no individual plaques could be identified, a faint lysis zone across the drop area was considered to be 10 plaques. The efficiency of plating was measured by comparing plaque assay results for control bacteria and those for bacteria containing the defense system and/or the defense system with a candidate anti-defense gene.

### Infection in liquid culture

Overnight cultures of bacteria harboring the defense system, negative control lacking the system, and stains with the defense system and candidate anti-defense gene pair were diluted 1:100 in MMB medium. 1 mM IPTG was added to induce expression of candidate anti-defense proteins. Cells were incubated at 37 °C while shaking at 200 rpm until early log phase (OD₆₀₀ of 0.3), and infected at a MOI of 0.01 in a 96-well plate containing 20 μL of phage lysate. Plates were incubated at 30 °C with shaking in a TECAN Infinite200 plate reader and OD₆₀₀ was followed with measurement every 10 min.

### NAD^+^ detection assay

Overnight cultures of bacteria harboring the Thoeris defense system, negative control cells lacking Thoeris and cells expressing Thoeris and candidate anti-defense genes were diluted 1:100 in MMB medium. 1mM IPTG was added to induce expression of candidate anti-defense proteins. Cells were incubated at 37 °C while shaking at 200 rpm until early log phase (OD₆₀₀ of 0.3), and infected at an MOI of 10 in a 96-well plate containing 20 μL of phage lysate. Plates were incubated at 30°C with shaking in a TECAN Infinite200 plate reader. 15 µL of cells were taken before phage infection (t=0) and at 30 and 60 minutes post infection. Cells were mixed with 20 µL of 100% ethanol and kept at -20 °C.

For the NAD^+^ detection assay, samples were diluted 1:5 in 100 mM sodium phosphate buffer (pH 7.5). Then 5 µL of the samples were mixed with 5 µL of NAD/NADH-Glo™ detection reagent (Promega corp.) in a Nunc^®^ 384 well polystyrene plate (Sigma p5991-1CS). Plates were incubated at 25°C in a TECAN Infinite200 plate reader and luminescence was followed with a measurement every 10 min. The reaction rate was calculated from the linear part of the initial reaction. Concentration values were determined with a calibration curve built with NAD^+^ samples of known concentrations.

### Protein pulldown assays

Proteins were tagged with C-terminal Twin-Strep tags. Each protein was expressed in a separate strain and lysates were then mixed for pulldowns, as described below.

Overnight cultures were diluted 1:100 in 50 mL MMB and grown for ∼1.5 h to OD_600_ ∼ 0.3 at 37 °C, shaking 200 rpm. Then, the appropriate inducer was added, and cells were incubated at 37 °C for an additional hour. Cells were then centrifuged at 3,900 *g* for 10 min. Supernatant was discarded and pellets were frozen at −80 °C.

To pull down the proteins, 1 mL of Strep-Tactin wash buffer (IBA catalogue no. 2-1003-100) supplemented with 4 mg mL^−1^ lysozyme was added to each pellet and incubated for 10 min at room temperature until thawed and then resuspended. Tubes were then transferred to ice, and the resuspended cells transferred to a FastPrep Lysing Matrix B in a 2 mL tube (MP Biomedicals catalogue no. 116911100). Samples were lysed using a FastPrep bead beater for two rounds of 40 s at 6 m s^−1^. Tubes were centrifuged for 15 min at 15,000 *g*. Per each pellet, 20 µl of MagStrep ‘Type 3’ XT beads (IBA catalogue no. 2-4090-002) were washed in 200 µl wash buffer (IBA catalogue no. 2-1003-100), and the lysed cell supernatant was mixed with the beads and incubated for 60 min, rotating at 4 °C. The sample mixed with the beads was comprised of equal parts (400 µL) of lysate for each protein tested, or in the case only one protein was tested, 400 µL of wash buffer was added to reach a total of 800 µL.

The beads were then pelleted on a magnet, washed twice with wash buffer, and purified protein was eluted from the beads in 40 µl of BXT elution buffer (IBA catalogue no. 2-1042-025). 30 µL of samples were mixed with 10 µL of 4× Bolt LDS Sample Buffer (ThermoFisher catalogue no. B0008) and dithiothreitol (DTT) to a final concentration of 1 mM. Samples were incubated at 75 °C for 5 min and then loaded to a NuPAGE 4% to 12%, Bis-Tris, 1.0 mm, Mini Protein Gel, 12 well (ThermoFisher catalogue no. NP0322PK2) in 20× Bolt MES SDS Running Buffer (ThermoFisher catalogue no. B0002) and run at 160 V. Gels were shaken with InstantBlue Coomassie Protein Stain (ISB1L) (ab119211) for 1 h, followed by another hour in water.

### Preparation of filtered cell lysates for NADase enzymatic assay

For generating filtered cell lysates, we grew *B. subtilis* BEST7003 cells co-expressing the candidate anti-defense proteins and the Thoeris system from *B. cereus* MSX-D12 in which ThsA was inactivated (ThsB + ThsA N112A) as described previously^28^. The anti-ThsB binder was integrated in the *thrC* locus and expressed from an inducible Phspank promoter^28^. The Thoeris system was integrated in the *amyE* locus and expressed from its native promoter. Controls included cells expressing the ThsB + ThsA(N112A) Thoeris system with GFP and cells that do not have Thoeris. All cultures were grown overnight and then diluted 1:100 in 250 mL MMB medium supplemented with 1 mM IPTG and grown at 37 °C, 200 rpm shaking for 120 min followed by incubation and shaking at 25 °C, 200 rpm until reaching an OD_600_ of 0.3. At this point, a sample of 50 mL was taken as the uninfected sample (time: 0 min), and the SBSphiJ phage was added to the remaining culture at an MOI of 10. Flasks were incubated at 25 °C with shaking (200 rpm), for the duration of the experiment. Samples of 50 mL were collected 105 minutes post infection. Immediately after sample removal the sample tubes were centrifuged at 4 °C for 10 min at 3,200 *g* to pellet the cells. The supernatant was discarded, and the pellet was flash frozen and stored at −80 °C. To extract cell metabolites from frozen pellets, 600 μL of 100 mM sodium phosphate buffer (pH 7.5) was added to each pellet. Samples were transferred to FastPrep Lysing Matrix B in a 2 mL tube (MP Biomedicals, cat #116911100) and lysed at 4 °C using a FastPrep bead beater for two rounds of 40 s at 6 ms^−1^. Tubes were then centrifuged at 4 °C for 10 min at 15,000 *g*. The supernatant was then transferred to an Amicon Ultra-0.5 Centrifugal Filter Unit 3 kDa (Merck Millipore, no. UFC500396) and centrifuged for 45 min at 4 °C, 12,000 *g*. Filtered lysates were taken for *in vitro* ThsA-based NADase activity assay.

### ThsA NADase enzymatic assay

The ThsA protein was expressed and purified as previously described^27^, and ThsA-based NADase activity assay for the detection of 3’cADPR was carried out as described previously^28^. NADase reaction was carried out in black 96-well half-area plates (Corning, cat #3694). In each reaction microwell, purified ThsA protein was added to cell lysate, or to 100 mM sodium phosphate buffer pH 8.0. A 5 μL volume of 5 mM nicotinamide 1,N6-ethenoadenine dinucleotide (εNAD^+^, Sigma, cat #N2630) solution was added to each well immediately before the beginning of measurements, resulting in a final concentration of 100 nM ThsA protein in a 50 μL final volume reaction. Plates were incubated inside a Tecan Infinite M200 plate reader at 25°C, and measurements were taken at 300 nm excitation wavelength and 410 nm emission wavelength. The reaction rate was calculated from the linear part of the initial reaction.

### Protein expression and purification

The ThsA protein was expressed under the control of the T7 promoter together with a C-terminal Twin-Strep tag for subsequent purification, as previously described^27^.

The gene encoding SLOG binder 2 was cloned into the pET28-bdSUMO expression vector, generating a construct with an N-terminal His-SUMO tag, as described in^62^. The plasmid was transformed into *E. coli* BL21 (DE3) cells. A 5-liter LB culture was grown at 37 °C until OD_600_ reached 0.6, at which point protein expression was induced with 0.2 mM IPTG. The culture was then incubated overnight at 15 °C.

Cells were harvested by centrifugation and the pellet was resuspended in lysis buffer consisting of PBS, 250 mM sucrose, and 1 mM DTT, supplemented with 200,000 U/100 mL lysozyme, 20 µg/mL DNase I, 1 mM MgCl₂, 1 mM PMSF, and a protease inhibitor cocktail. Cell lysis was performed using a cooled cell disrupter (Constant Systems).

The clarified lysate was incubated with 5 mL of pre-washed Ni-NTA agarose beads (Adar Biotech) for 1 hour at 4 °C. Following removal of unbound proteins, the beads were washed four times with 50 mL of lysis buffer.

To elute the target protein, the beads were incubated with 5 mL of lysis buffer supplemented with 0.2 mg bdSUMO protease for 2 hours at room temperature. The cleaved SLOG binder 2 was collected from the supernatant and further purified by ion exchange chromatography using a Capto HiRes Q 10/100 column (Cytiva), pre-equilibrated with 50 mM Tris-HCl, pH 8.0.

Elution was carried out using a linear NaCl gradient up to 1 M. SLOG binder 2 eluted at approximately 100 mM NaCl and was collected as the pure fraction. The purified protein was aliquoted, flash-frozen in liquid nitrogen, and stored at –80 °C.

### SPR Protein-Protein binding assay

Surface plasmon resonance (SPR) experiments were performed using Biacore S200 (Cytiva). The SLOG binder 2 was immobilized onto a Biacore sensor chip CM5 Series S (Cytiva) using Amine coupling kit (Cytiva), to final immobilization values of 430 RUs. Two-fold serial dilutions of purified ThsA protein, ranging from 1.5 to 25 nM were injected for a duration of 120 sec, and dissociation was observed for 1200 sec, at a flow rate of 30 µl/min, using PBST as a running buffer, at 25 °C. For regeneration between cycles, NaOH, 1 mM, was injected for 30 sec, at 10 µl/min. Sensograms were fitted using the Biacore Insight Evaluation Software (Cytiva, v 4.0.8.19879), using the two-state reaction model, resulting in KD of ∼60 nM. The experiment was repeated three times with comparable results.

### Pooled candidate plasmid approach for DdmDE binders

Candidate DdmDE binders were ordered from Twist Bioscience as a ssDNA oligo pool with primer binder sites for PCR amplification and downstream Gibson assembly. The oligo pool was amplified for 14 cycles with the primers gaccgaattcaaaagatcttttaagaaggagatatacatatg and gcctggagatccttactcgagtttggatcc . The resulting PCR reaction was added to a NEBuilder HiFi DNA Assembly kit (NEB, cat # E5520S) reaction with a pBbA6c (addgene plasmid #35290) backbone (primers ggatccaaactcgagtaaggatctcca and atgtatatctccttcttaaaagatcttttgaattcggt). The resulting reaction was transformed into DH5α (NEB 5-alpha Competent *E. coli)* cells. Roughly 10,000 colonies were collected and mixed as a pooled library of candidate plasmid-containing cells.

50 µL of the DH5α library cells were mini prepped and the obtained plasmids were used to transform electro competent *E. coli* cells containing genomically integrated DdmDE or an empty cassette (control cells). Roughly 10,000 colonies were collected and mixed as a pooled library of candidate plasmids in DdmDE containing cells or control cells. For sequencing of the initial libraries, 50 µL of cells were mini prepped and the obtained plasmids were used as templates for 14 cycle PCR amplification of the candidate region (primer caaaagatcttttaagaaggagatatacatatg and ccttactcgagtttggatcc ). The resulting PCR reaction was sent to Plasmidsaurus for PCR premium sequencing and raw long reads were used for downstream analysis.

5 µL of DdmDE or control cell libraries were used to start overnight cultures with chloramphenicol (30 μg mL^−1^) and 1% glucose. The following day, cultures were diluted 1:100 in MMB medium. 1mM IPTG and 0.2% arabinose was added to induce expression of candidate anti-defense proteins and the DdmDE system. Cells were incubated at 30 °C while shaking at 200 rpm for 50 generations. The cultures were diluted and plated on LB plates with or without chloramphenicol (30 μg mL^−1^) and roughly 10,000 colonies were collected. As before, 50 µL of cells were mini prepped, and 14 cycle PCR amplifications of candidate regions of plasmids were sent to sequencing.

Raw, long reads were obtained for all sequencing data (Plasmidsaurus premium PCR sequencing). For all conditions, the total amount of reads per candidate were normalized to the total amount of reads per condition. Enrichment was calculated by dividing the normalized amount of reads per candidate at the end of the experiment to those measured at the beginning of the experiment in control cells. Candidates that showed >5-fold enrichment in all three replicated experiments were ordered for individual testing.

### Plasmid elimination assay in DdmDE-expressing *E. coli*

Candidate binders were ordered from Twist Bioscience as gene fragments with adaptor sequences for downstream Gibson assembly with the pBbA6c plasmid backbone. The resulting NEBuilder HiFi DNA Assembly reactions and the RFP expressing pBbA6c control plasmid were transformed into electro competent *E. coli* cells containing genomically integrated DdmDE. Bacteria harboring DdmDE and a candidate binder or control plasmid were used to start overnight cultures with chloramphenicol (30 μg mL^−1^) and 1% glucose. The following day, cultures were diluted 1:100 in MMB medium. 1mM IPTG and 0.2% arabinose was added to induce expression of candidate anti-defense proteins and the DdmDE system. Cells were incubated at 30°C while shaking at 200 rpm for 50 generations. The cultures were serially diluted in PBS and 10 µL of each dilution (10^0^-10^-7^) was spotted on LB agar plates, with or without chloramphenicol (30 μg mL^−1^) selection. Following incubation overnight at 30°C, plasmid retention was calculated as the ratio of the chloramphenicol-resistant CFU (i.e. plasmid-carrying cells) / total number of bacteria CFU.

### Plasmid elimination assay in *V. cholerae*

Plasmid stability assays were conducted in the 7th pandemic O1 El Tor *V. cholerae* strain A1552^58^, as previously described^16^. Briefly, overnight cultures grown with selection (Kanamycin 75µg/mL) were back-diluted 1:1250 in Lysogeny broth (LB-Miller; 10 g/L tryptone, 5 g/L yeast extract, 10 g/L NaCl; Carl Roth, Switzerland) and cultured at 30°C, 180rpm without selection for approximately 50 generations, with dilution into fresh media every 10 generations. Next, cultures were serially diluted in PBS and 5 µL of each dilution (10^0^-10^-7^) spotted in duplicate on LB agar plates, without and with selection. Following incubation overnight at 30°C, plasmid retention was calculated as the ratio of the kanamycin-resistant CFU (i.e. plasmid-carrying cells) / total number of bacteria.

### De novo binder design using RFdiffusion

Cropped AF2 models of defense proteins served as the target inputs for RFdiffusion. In the case of the SLOG receptor, the sequence of two cropped SLOG domains were joined with a 29-residue long poly G flexible linker and the resulting AF2-predicted model was used as the target. Approximately 10,000 designs were generated using the RFdiffusion model. The resulting backbone libraries underwent sequence design using ProteinMPNN, followed by FastRelax and AF2 initial guess approach ^33^. The resulting libraries were filtered based on AF2 predicted aligned error (PAE) < 10. Where not already present, a Methionine residue was added to the N-terminus of protein sequences ordered for experimentation.

### Partial diffusion to optimize binders using RFdiffusion

For the DdmDE PIWI and nuclease domain targets, which were hard to obtain high scoring binders against, the AF2 models of the highest-scoring design was used as the input for partial diffusion using the RFdiffusion model. The models were subjected to 20 noising time steps out of a total of 50 time steps in the noising schedule and subsequently denoised (“diffuser.partial_T” input values of 20). Approximately 10,000 partially diffused designs were generated for each target. The resulting library of backbones was sequence designed using ProteinMPNN after Rosetta FastRelax, followed by AF2 initial guess approach ^33^. The resulting libraries were filtered based on AF2 PAE < 10.

### Prediction of protein structures and interactions

Where stated, AlphaFold3^63^ was used to generate structural models for proteins and protein complexes. The chain pair interface-predicted TM (ipTM) score is noted where applicable.

In Fig. 1b the binder is folded with two SLOG domains connected with a 29-residue long poly G linker. In Fig. 3a, due to its size, the Avs1 B tetramer was modeled by joining and aligning two AF3 models. The displayed binder is folded with a cropped part of the TPR domain (residues 621-1117 of the Avs1B protein) and then structurally aligned to the full Avs1B model. The ipTM score is derived from folding the binder with the full Avs1B protein. In Fig. 5e, binders 1-4 were each folded with the relevant protein (DdmD or DdmE). Binder 5 was designed to interfere with the assembly of the DdmDE complex, and was hence folded with the cropped section of the DdmE protein (residues 286-412) used as input for RFdiffusion and then aligned to the DdmDE structure. In Fig. S4, the binders were folded with two SLOG domains connected with a 29-residue long poly G linker and ipTM scores were obtained. The models were then aligned to PDB structure ID 7UXS and the binder was shown in place.

### Phage competition

Bacteria expressing the appropriate defense systems or control cells with no defense were grown to OD ∼ 0.3 at 37°C in MMB media. The WT SBSphiJ phage and the synthetic binder-containing phages were mixed to 1:1 PFU ratio, and the mix was used to infect the bacterial cultures with an MOI of 0.1. The infection proceeded at 30 °C for five hours, and then the cultures were centrifugated 15 min at 3200 *g* and filtered to recover the phages. For the next two days, a similar infection was performed using the phages from the previous day to infect a new bacterial culture at low MOI. After three passages of the mixed phages, a PCR amplification of the region encoding the knock in genes was performed and visualized on an agarose gel. The intensity of the DNA band corresponding to either the WT or knock in fragment was used as a proxy for phage abundance.

## Acknowledgements

We thank members of the Sorek lab for constructive criticism during this study. R.S. was supported, in part, by the European Research Council (grant ERC-AdG GA 101018520), the Israel Science Foundation (MAPATS grant 2720/22), the Minerva Foundation with funding from the Federal German Ministry for Education and Research, a research grant from the Estate of Hermine Miller, the Institute for Environmental Sustainability (IES) and the Center for Immunotherapy at the Weizmann Institute of Science, and the Knell Family Center for Microbiology. M.B. acknowledges support from the Swiss National Science Foundation (project grant 320030-227514) and EPFL intramural funding.

## Supplemental figures

**Figure S1.**
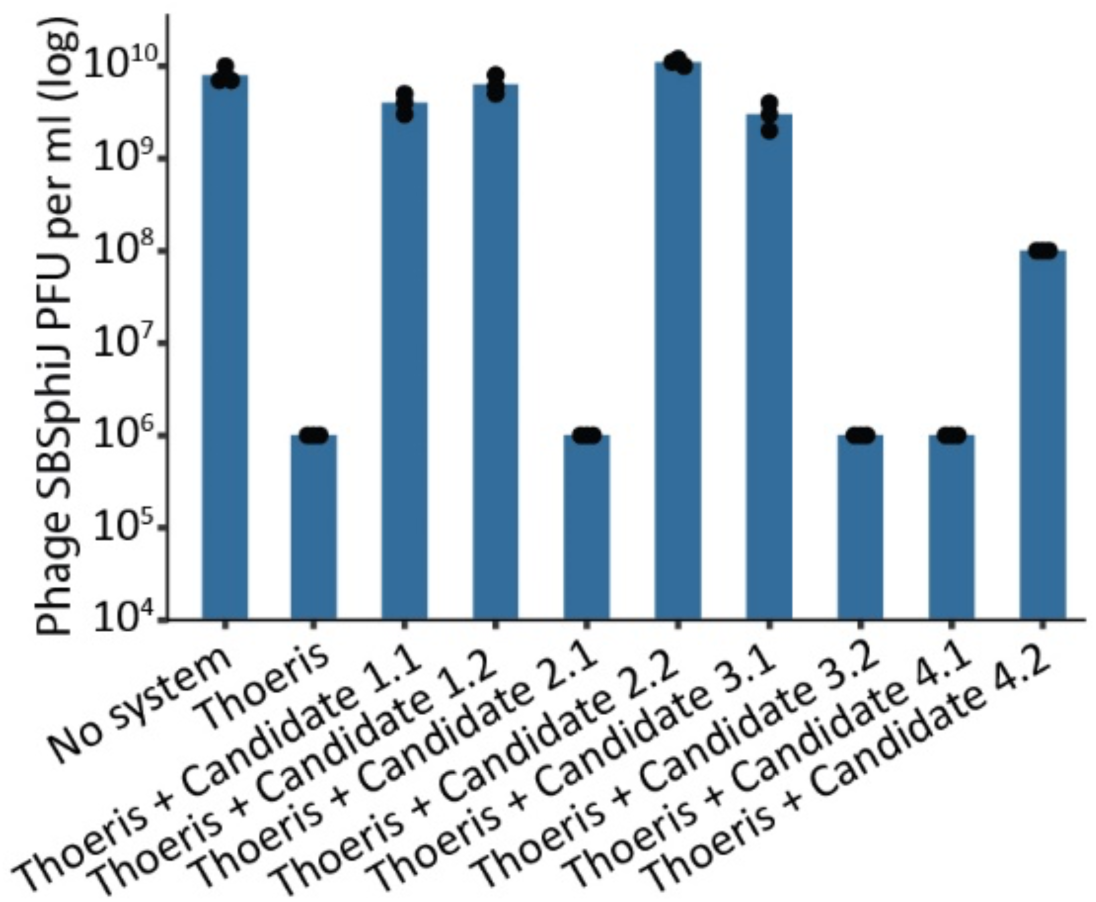
Anti-defense activity of individual ThsA binders from verified protein pairs. Two-gene constructs shown to inhibit Thoeris defense in liquid culture experiments were separated and tested as individual genes in plaque assays. Shown are efficiency of plating (EOP) data for phage SBSphiJ infecting the control *B. subtilis* BEST7003 strain (no system), *B. subtilis* BEST7003 with Thoeris cloned from *B. cereus* MSX-D12, or cells co-expressing Thoeris and single candidate binders. Data represent plaque-forming units (PFU) per milliliter. Bar graphs represent average of three independent replicates, with individual data points overlaid.

**Figure S2.**
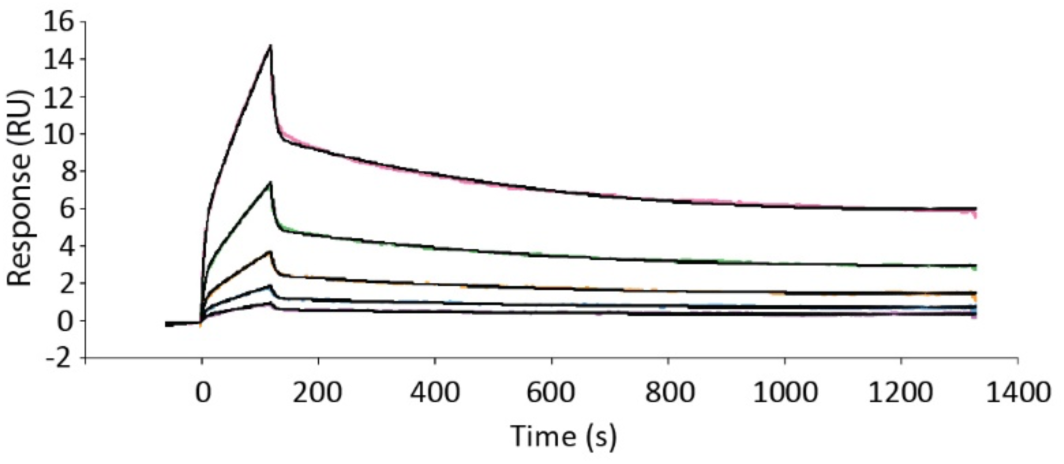
Affinity of SLOG binder 2 to ThsA, as measured by SPR. Increasing concentrations (1.5 nM to 25 nM with two-fold intervals) of ThsA protein were injected over immobilized SLOG binder 2, using PBST buffer, and the response (RU) was followed over time (colored lines). Data was fitted using a two-state reaction model (black lines), resulting in a dissociation constant (Kd) of ∼60 nM.

**Figure S3.**
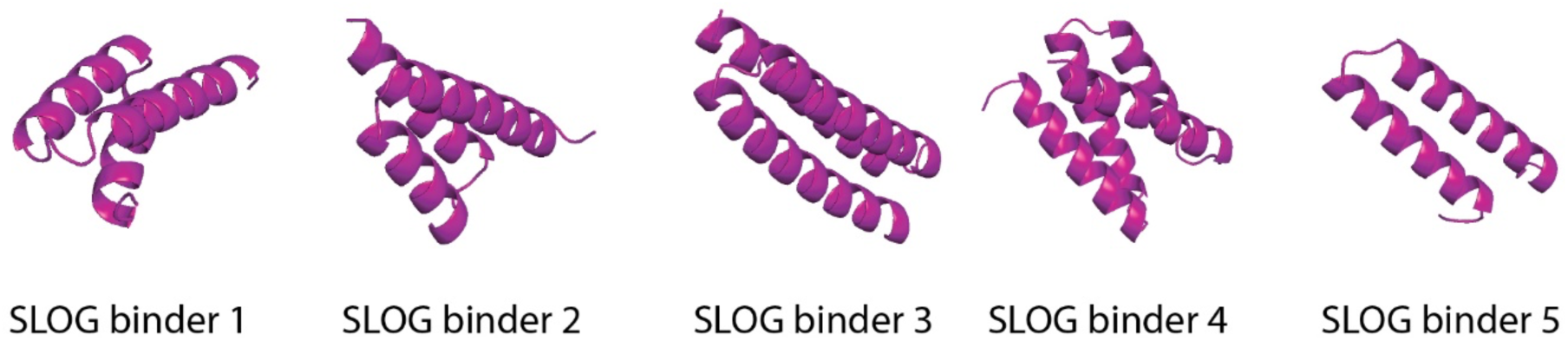
**AF3-predicted structure models of verified synthetic SLOG binders.**

**Figure S4.**
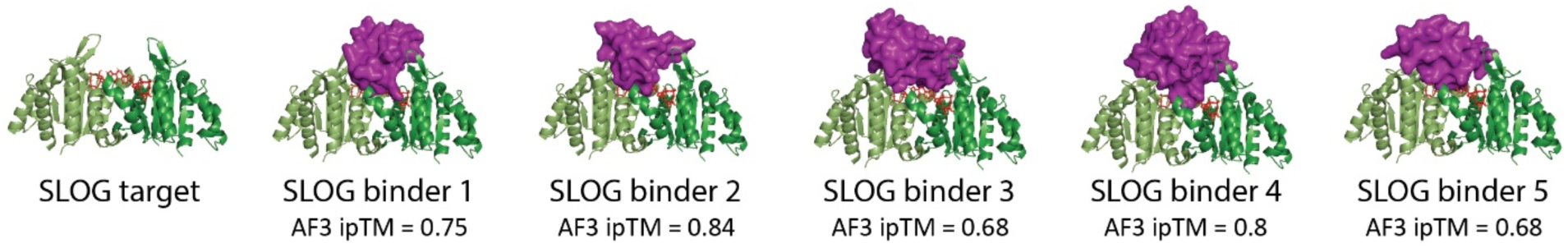
AF3-predicted structures of verified SLOG binders modeled with the SLOG domain of ThsA. The unbound SLOG domain in complex with 3’cADPR is from PDB ID 7UXS. ipTM scores are of AF3 predictions of binders folded with SLOG target (two cropped SLOG domains joined with a poly glycine linker).

**Figure S5.**
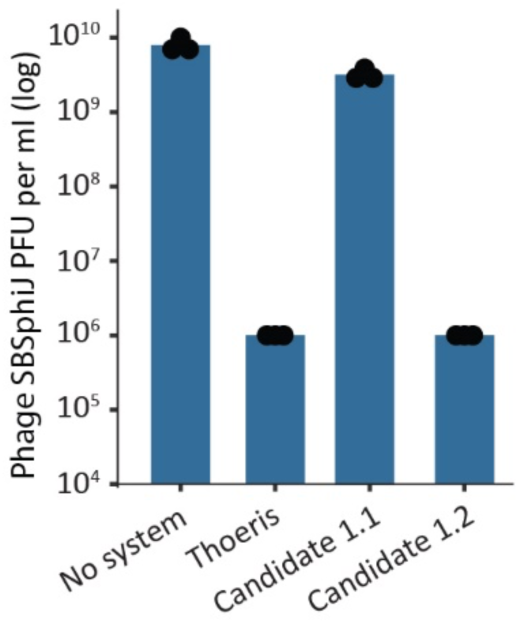
Anti-defense activity of individual designed ThsB binders from a verified protein pair. The two-protein constructs shown to inhibit Thoeris defense in liquid culture experiments was separated and tested as individual proteins in plaque assays. Shown are efficiency of plating (EOP) data for phage SBSphiJ infecting the control *B. subtilis* BEST7003 strain (no system), *B. subtilis* BEST7003 with Thoeris cloned from *B. cereus* MSX-D12, or cells co-expressing Thoeris and single candidate binders. Data represent PFU per milliliter. Bar graphs represent average of three independent replicates, with individual data points overlaid.

**Figure S6.**
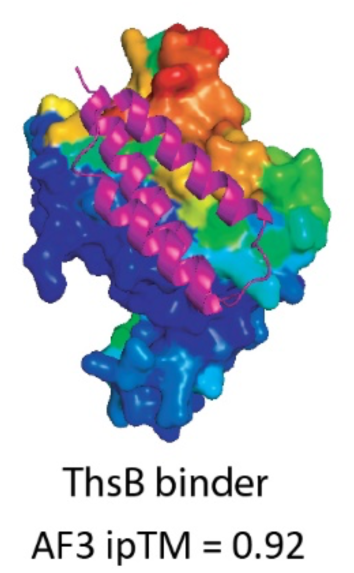
A predicted protein-protein interaction interface in ThsB. (a) ScanNet^64^ protein binding site prediction. Low (blue) to high (red) probability of protein-protein interaction. AF3-predicted structure of the ThsB binder is modeled in place.

**Figure S7.**
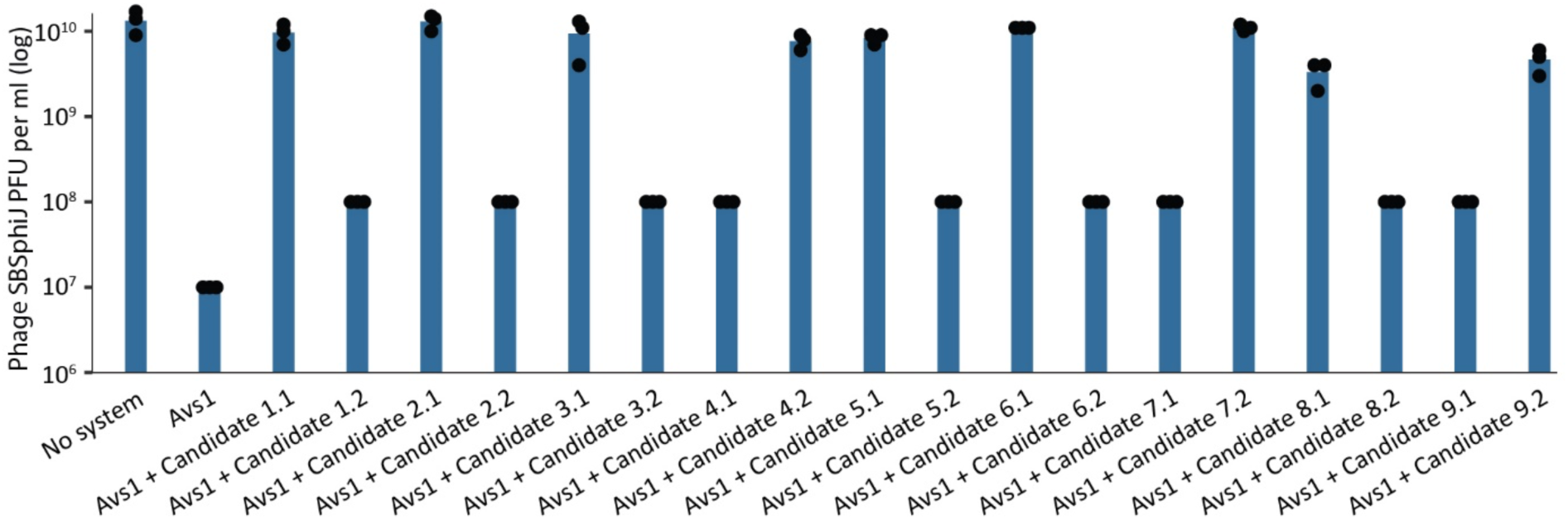
Anti-defense activity of individual designed Avs1 binders from verified protein pairs. Two-protein constructs shown to inhibit Avs1 defense in liquid culture experiments were separated and tested as individual proteins in plaque assays. Shown are efficiency of plating (EOP) data for phage SBSphiJ infecting the control *B. subtilis* BEST7003 strain (no system), *B. subtilis* BEST7003 with Avs1 cloned from *B. cereus* VD102, or cells co-expressing Avs1 and single candidate binders. Data represent PFU per milliliter. Bar graphs represent average of three independent replicates, with individual data points overlaid.

**Figure S8.**
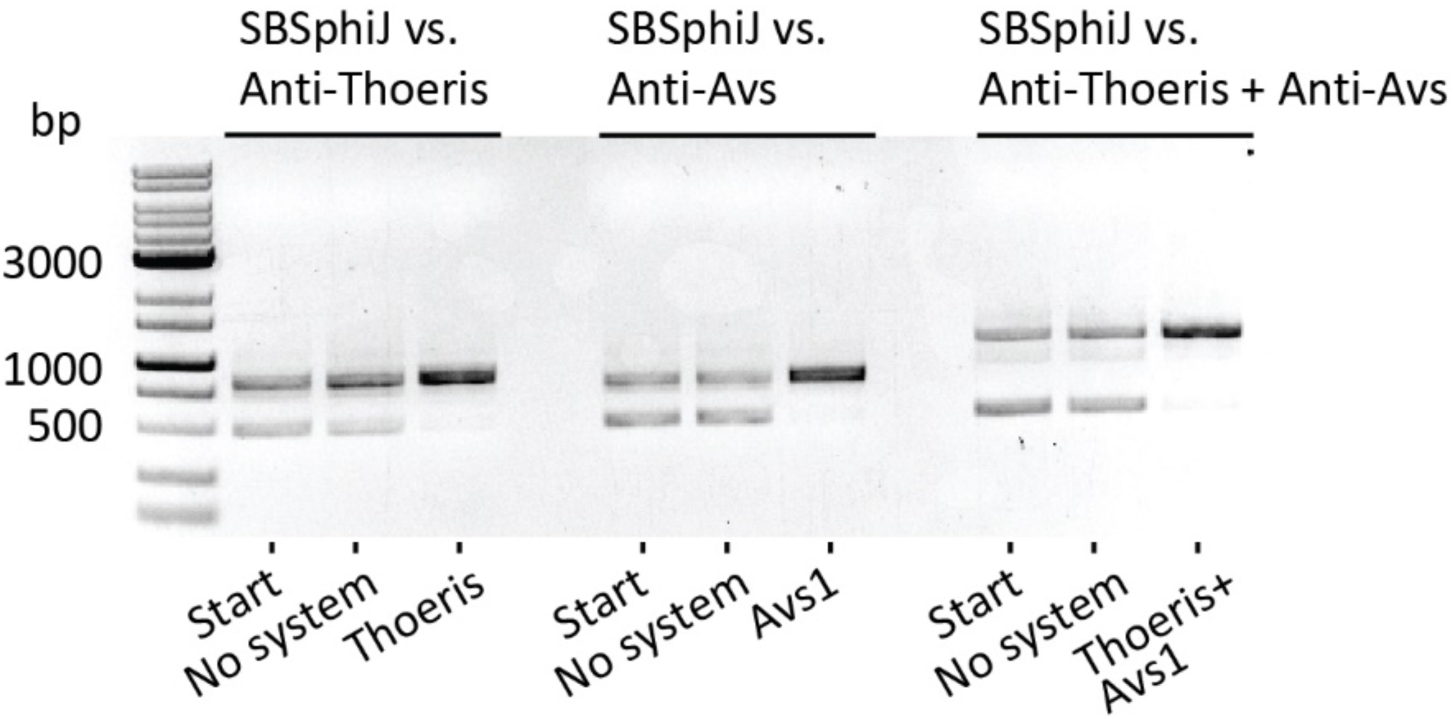
Synthetic proteins do not inflict fitness cost on engineered phages. Wild type phage SBSphiJ and SBSphiJ knocked in with synthetic inhibitors of defense systems were mixed to a 1:1 ratio. The mix was then used to infect control cells (no system) or cells expressing the appropriate system in liquid culture. Shown are the results of a PCR amplification of the knock in region of the mixed phage population before competition (start) and after 3 rounds of infection. The lower band corresponds to the wild type, non engineered phage; the upper band corresponds to the knocked-in DNA fragment, used here as a proxy for the corresponding phage abundance. A DNA ladder in base pair (bp) is presented on the left.

**Figure S9.**
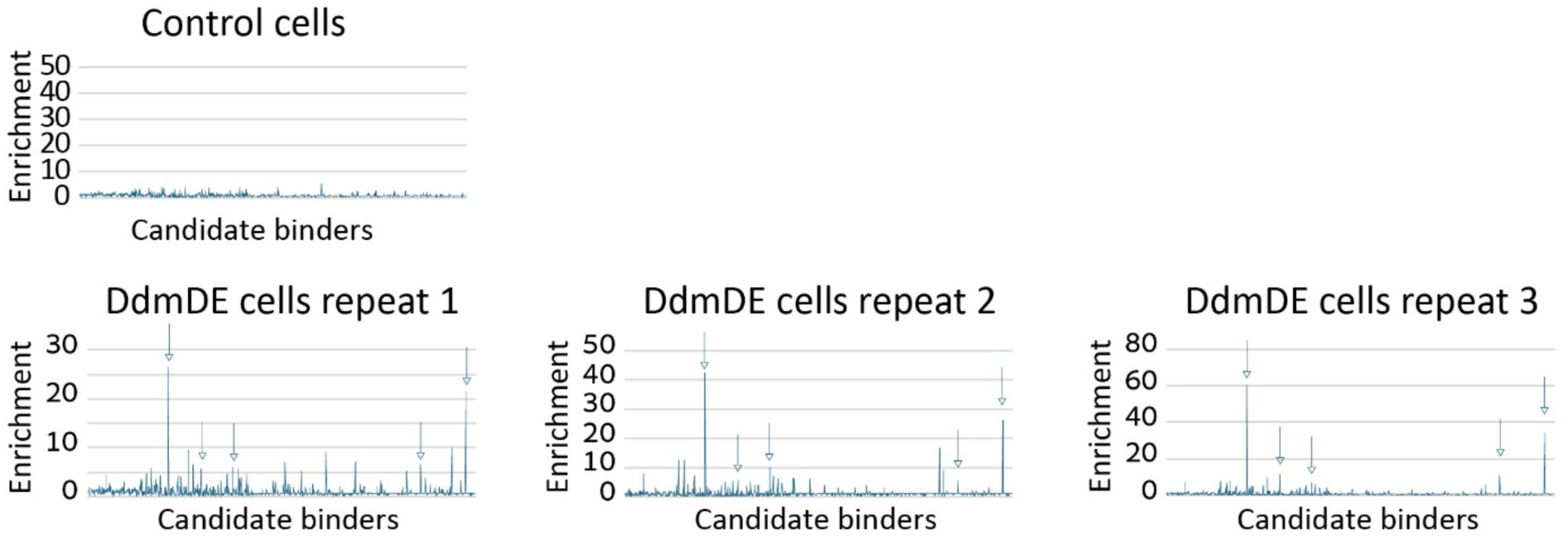
Three replicates of the anti-DdmDE plasmid enrichment experiment. Enrichment was calculated by dividing reads per candidate in the initial library by the reads per candidate at the end point of the experiment. Read counts were normalized to total reads per sample. Top, no enrichment of candidate plasmids in control cells. Bottom, enrichment of candidate plasmids in DdmDE expressing cells in three replicates. Candidates that showed >5-fold enrichment in all three replicates are indicated with arrows. Data for control and repeat 3 are presented also in Figure 5.

## Supplemental tables

**Table S1.** Validated de-novo synthesized defense inhibitors.

**Table S2.** Phages engineered in this study

## Notes

### Competing Interest Statement

R.S. is a scientific cofounder and advisor of BiomX and Ecophage. The other authors declare no competing interests.

## References

1. Georjon, H. & Bernheim, A. The highly diverse antiphage defence systems of bacteria. Nat. Rev. Microbiol. 21, 686–700 (2023).

2. Doron, S. et al. Systematic discovery of antiphage defense systems in the microbial pangenome. Science 359, eaar4120 (2018).

3. Millman, A. et al. An expanded arsenal of immune systems that protect bacteria from phages. Cell Host Microbe 30, 1556–1569.e5 (2022).

4. Gao, L. et al. Diverse enzymatic activities mediate antiviral immunity in prokaryotes. Science 369, 1077–1084 (2020).

5. Rousset, F. et al. Phages and their satellites encode hotspots of antiviral systems. Cell Host Microbe 30, 740–753.e5 (2022).

6. Vassallo, C. N., Doering, C. R., Littlehale, M. L., Teodoro, G. I. C. & Laub, M. T. A functional selection reveals previously undetected anti-phage defence systems in the E. coli pangenome. Nat. Microbiol. 7, 1568–1579 (2022).

7. Tesson, F. et al. Systematic and quantitative view of the antiviral arsenal of prokaryotes. Nat. Commun. 13, 2561 (2022).

8. Hochhauser, D., Millman, A. & Sorek, R. The defense island repertoire of the Escherichia coli pan-genome. PLOS Genet. 19, e1010694 (2023).

9. Bernheim, A. & Sorek, R. The pan-immune system of bacteria: antiviral defence as a community resource. Nat. Rev. Microbiol. 18, 113–119 (2020).

10. Bleriot, I., et al. Improving phage therapy by evasion of phage resistance mechanisms. JAC-Antimicrob. Resist. 6, dlae017 (2023).

11. Egido, J. E., Costa, A. R., Aparicio-Maldonado, C., Haas, P.-J. & Brouns, S. J. J. Mechanisms and clinical importance of bacteriophage resistance. FEMS Microbiol. Rev. 46, fuab048 (2022).

12. Skurnik, M. et al. Phage therapy. Nat. Rev. Methods Primer 5, 9 (2025).

13. Sarker, S. A. & Brüssow, H. From bench to bed and back again: phage therapy of childhood *Escherichia coli* diarrhea. Ann. N. Y. Acad. Sci. 1372, 42–52 (2016).

14. Dedrick, R. M. et al. Engineered bacteriophages for treatment of a patient with a disseminated drug-resistant Mycobacterium abscessus. Nat. Med. 25, 730–733 (2019).

15. Oliveira, P. H., Touchon, M. & Rocha, E. P. C. The interplay of restriction-modification systems with mobile genetic elements and their prokaryotic hosts. Nucleic Acids Res. 42, 10618–10631 (2014).

16. Jaskólska, M., Adams, D. W. & Blokesch, M. Two defence systems eliminate plasmids from seventh pandemic Vibrio cholerae. Nature 604, 323–329 (2022).

17. Zaremba, M. et al. Short prokaryotic Argonautes provide defence against incoming mobile genetic elements through NAD+ depletion. Nat. Microbiol. 7, 1857–1869 (2022).

18. Koopal, B. et al. Short prokaryotic Argonaute systems trigger cell death upon detection of invading DNA. Cell 185, 1471–1486.e19 (2022).

19. Baker, B. J., Hyde, E. & Leão, P. Nature should be the model for microbial sciences. J. Bacteriol. 206, e00228–24 (2024).

20. Deep, A. et al. The SMC-family Wadjet complex protects bacteria from plasmid transformation by recognition and cleavage of closed-circular DNA. Mol. Cell 82, 4145–4159.e7 (2022).

21. Robins, W. P., Meader, B. T., Toska, J. & Mekalanos, J. J. DdmABC-dependent death triggered by viral palindromic DNA sequences. Cell Rep. 43, 114450 (2024).

22. Loeff, L. et al. Molecular mechanism of plasmid elimination by the DdmDE defense system. Science 385, 188–194 (2024).

23. Listov, D., Goverde, C. A., Correia, B. E. & Fleishman, S. J. Opportunities and challenges in design and optimization of protein function. Nat. Rev. Mol. Cell Biol. 25, 639–653 (2024).

24. Watson, J. L. et al. De novo design of protein structure and function with RFdiffusion. Nature 620, 1089–1100 (2023).

25. Gainza, P. et al. De novo design of protein interactions with learned surface fingerprints. Nature 617, 176–184 (2023).

26. Pacesa, M., et al. BindCraft: one-shot design of functional protein binders. bioRxiv (2024) doi:10.1101/2024.09.30.615802.

27. Ofir, G. et al. Antiviral activity of bacterial TIR domains via immune signalling molecules. Nature 600, 116–120 (2021).

28. Leavitt, A. et al. Viruses inhibit TIR gcADPR signalling to overcome bacterial defence. Nature 611, 326–331 (2022).

29. Manik, M. K. et al. Cyclic ADP ribose isomers: Production, chemical structures, and immune signaling. Science 377, eadc8969 (2022).

30. Tamulaitiene, G. et al. Activation of Thoeris antiviral system via SIR2 effector filament assembly. Nature 627, 431–436 (2024).

31. Ka, D., Oh, H., Park, E., Kim, J.-H. & Bae, E. Structural and functional evidence of bacterial antiphage protection by Thoeris defense system via NAD+ degradation. Nat. Commun. 11, 2816 (2020).

32. Dauparas, J. et al. Robust deep learning–based protein sequence design using ProteinMPNN. Science 378, 49–56 (2022).

33. Bennett, N. R. et al. Improving de novo protein binder design with deep learning. Nat. Commun. 14, 2625 (2023).

34. Schmidt, T. G. M. et al. Development of the Twin-Strep-tag® and its application for purification of recombinant proteins from cell culture supernatants. Protein Expr. Purif. 92, 54–61 (2013).

35. Gao, L. A. et al. Prokaryotic innate immunity through pattern recognition of conserved viral proteins. Science 377, eabm4096 (2022).

36. Bernheim, A., Cury, J. & Poirier, E. Z. The immune modules conserved across the tree of life: Towards a definition of ancestral immunity. PLOS Biol. 22, e3002717 (2024).

37. Kibby, E. M. et al. Bacterial NLR-related proteins protect against phage. Cell 186, 2410–2424.e18 (2023).

38. Chou, W.-C., Jha, S., Linhoff, M. W. & Ting, J. P.-Y. The NLR gene family: from discovery to present day. Nat. Rev. Immunol. 23, 635–654 (2023).

39. Béchon, N. et al. Diversification of molecular pattern recognition in bacterial NLR-like proteins. Nat. Commun. 15, 9860 (2024).

40. Kanungo, S., Azman, A. S., Ramamurthy, T., Deen, J. & Dutta, S. Cholera. The Lancet 3GG, 1429–1440 (2022).

41. Newland, J. W., Voll, M. J. & McNicol, L. A. Serology and plasmid carriage in Vibrio cholerae. Can. J. Microbiol. 30, 1149–1156 (1984).

42. Amaro, C., Aznar, R., Garay, E. & Alcaide, E. R plasmids in environmental Vibrio cholerae non-O1 strains. Appl. Environ. Microbiol. 54, 2771–2776 (1988).

43. Weill, F.-X. et al. Genomic history of the seventh pandemic of cholera in Africa. Science 358, 785–789 (2017).

44. Bravo, J. P. K., Ramos, D. A., Fregoso Ocampo, R., Ingram, C. & Taylor, D. W. Plasmid targeting and destruction by the DdmDE bacterial defence system. Nature 630, 961–967 (2024).

45. Yang, X.-Y., Shen, Z., Wang, C., Nakanishi, K. & Fu, T.-M. DdmDE eliminates plasmid invasion by DNA-guided DNA targeting. Cell 187, 5253–5266.e16 (2024).

46. Chauthaiwale, V. M., Therwath, A. & Deshpande, V. V. Bacteriophage lambda as a cloning vector. Microbiol. Rev. 56, 577–591 (1992).

47. Liu, Y., Huang, H., Wang, H. & Zhang, Y. A novel approach for T7 bacteriophage genome integration of exogenous DNA. J. Biol. Eng. 14, 2 (2020).

48. Costa, A. R. et al. Accumulation of defense systems in phage-resistant strains of Pseudomonas aeruginosa. Sci. Adv. 10, eadj0341.

49. Baca, C. F. & Marraffini, L. A. Nucleic acid recognition during prokaryotic immunity. Mol. Cell 85, 309–322 (2025).

50. Riley, L. A. & Guss, A. M. Approaches to genetic tool development for rapid domestication of non-model microorganisms. Biotechnol. Biofuels 14, 30 (2021).

51. Hopwood, D. A., Wright, H. M., Bibb, M. J. & Cohen, S. N. Genetic recombination through protoplast fusion in Streptomyces. Nature 268, 171–174 (1977).

52. Itaya, M. et al. Stable and efficient delivery of DNA to *Bacillus subtilis* ( *natto* ) using pLS20 conjugational transfer plasmids. FEMS Microbiol. Lett. 366, (2019).

53. Monk, I. R. Genetic manipulation of Staphylococci—breaking through the barrier. Front. Cell. Infect. Microbiol. 2, (2012).

54. Monk, I. R., Shah, I. M., Xu, M., Tan, M.-W. & Foster, T. J. Transforming the Untransformable: Application of Direct Transformation To Manipulate Genetically Staphylococcus aureus and Staphylococcus epidermidis. mBio 3, e00277–11 (2012).

55. Islam, Z. F. et al. Two Chloroflexi classes independently evolved the ability to persist on atmospheric hydrogen and carbon monoxide. ISME J. 13, 1801–1813 (2019).

56. Aune, T. E. V. & Aachmann, F. L. Methodologies to increase the transformation efficiencies and the range of bacteria that can be transformed. Appl. Microbiol. Biotechnol. 85, 1301–1313 (2010).

57. Yirmiya, E. et al. Structure-guided discovery of viral proteins that inhibit host immunity. Cell 188, 1681–1692.e17 (2025).

58. Yildiz, F. H. & Schoolnik, G. K. Role of *rpoS* in Stress Survival and Virulence of *Vibrio cholerae*. J. Bacteriol. 180, 773–784 (1998).

59. Matthey, N., Drebes Dörr, N. C. & Blokesch, M. Long-Read-Based Genome Sequences of Pandemic and Environmental Vibrio cholerae Strains. Microbiol. Resour. Announc. 7, 10.1128/mra.01574-18 (2018).

60. Osterman, I. et al. Phages reconstitute NAD+ to counter bacterial immunity. Nature 634, 1160–1167 (2024).

61. Mazzocco, A., Waddell, T. E., Lingohr, E. & Johnson, R. P. Enumeration of Bacteriophages Using the Small Drop Plaque Assay System. Methods Mol. Biol. 501, 81–85 (2009).

62. Zahradník, J. et al. Flexible regions govern promiscuous binding of IL-24 to receptors IL-20R1 and IL-22R1. FEBS J. 286, 3858–3873 (2019).

63. Abramson, J. et al. Accurate structure prediction of biomolecular interactions with AlphaFold 3. Nature 630, 493–500 (2024).

64. Tubiana, J., Schneidman-Duhovny, D. & Wolfson, H. J. ScanNet: an interpretable geometric deep learning model for structure-based protein binding site prediction. Nat. Methods 19, 730–739 (2022).

